# Altered food habits? Understanding the feeding preference of free-ranging Grey langur (*Semnopithecus entellus*) within an urban settlement

**DOI:** 10.1101/2020.10.22.350926

**Authors:** Dishari Dasgupta, Arnab Banerjee, Rikita Karar, Debolina Banerjee, Shohini Mitra, Purnendu Sardar, Srijita Karmakar, Aparajita Bhattacharya, Swastika Ghosh, Pritha Bhattacharjee, Manabi Paul

**Author notes:** Department of Environmental Science, University of Calcutta, 35, Ballygunge Circular Road, Kolkata-700019, India., *email: (Manabi Paul).

## Abstract

Urbanization affects concurrent human-animal movements as a result of altered resource availability and land use pattern, which leads to considerable ecological consequences. While some animals find themselves adrift, homeless with the uncertainty of resources resulting from the urban encroachment, few of them manage to survive by altering their natural behavioural patterns, and co-exist with humans. Folivorous colobines, such as grey langur, whose feeding repertoire largely consists of plant parts, tend to be more attuned to the urban high-calorie food sources to attain maximum fitness benefits within the concrete jungle having an insignificant green cover. However, such a mismatch between their generalized feeding behaviour and specialized gut physiology reminds us of the Liem’s paradox and demands considerable scientific attention which could tell us the story behind colobines’ successful co-existence within human settlements. Besides understanding their population dynamics, the effective management of these urbanized, free-ranging, non-human primate populations also depends on their altered feeding preferences, altogether which could lead us to the development of an ecologically sound urban ecosystem. Here, we have used a field-based experimental set up which allows langurs to choose between natural and urban food options, being independent of any inter-specific conflicts over resources due to food scarcity. The multinomial logit model reveals the choice-based decision making of these free-ranging grey langurs in an urban settlement of West Bengal, India, where they have not only learned to approach the human-provisioned urban food items but also shown a keen interest in it. While urbanization imposes tremendous survival challenges to these animals, it also opens up for various alternative options for human-animal co-existence which is reflected in this study, and could guide us for the establishment of a sustainable urban ecosystem in the future.

**HIGHLIGHTS:**

- The feeding repertoire of free-ranging grey langurs at Dakshineswar largely consists of urban food items in contrast to the langurs of Nangi, and Nalpur who mostly depend on natural food sources.
- High human-langur interactions together with the scarcity of natural plant-based food sources could be considered as an intriguing driving force behind langurs’ altered feeding habits in Dakhineswar.
- The field-based experimental set up allows free-ranging langurs to choose between natural and urban food options in an urban settlement like Dakhineswar.
- Urban food items outperformed natural food items as the most chosen one, indicating langurs’ altered feeding preferences which facilitate their successful co-existence within an urban ecosystem.

---

The global urban human population is set to reach the five billion mark by 2028 (ONU, 2018), facilitating urban sprawling and subsequently contributing to natural habitat loss worldwide at an unprecedented rate. This is expected to affect a large number of animals whose habitat ranges overlap with urban areas (He et al., 2014; Martinuzzi et al., 2015; Mcdonald et al., 2008; Murray & St. Clair, 2015). Habitat fragmentation and encroachment due to such urban expansion, which is often irreversible, has forced many homeless animals to live in close proximity to humans (Bateman & Fleming, 2012), giving rise to frequent human-animal conflict (Devi & Saikia, 2008; Omondi et.al.,2004; Messmer 2000; Woodruffe et.al.,2005). At the same time, some of these animals have also shown considerable behavioural adaptations like altered nesting or denning habits, vocalization, migratory activities, mating and breeding patterns, feeding behaviour (Able & Belthoff, 1998; Estes & Mannan, 2003; Kettlewell, 1961; Lowry et al., 2013; Slabbekoorn & Peet, 2003; Swedell et al., 2011) together with life history modification to survive amidst anthropogenic stress. Such anthropogenic stress often creates unpredictable selection pressure on these urban animals, leading to a sharp decline in species richness and composition within an urban ecosystem (Erinjery et al., 2017; Fuentes, 2012; Kale et al., 2012; H. N. Kumara & Singh, 2004; Paul et al., 2016; Singh & Raghunatha Rao, 2004; Vitousek et al., 1997). However, despite significant loss of biodiversity, urban expansion offers various high-calorie resource options to the generalist species who have higher dietary as well as foraging plasticity, and therefore, can adjust more readily to the altered habitat in contrast to the specialists (Fisher & Owens, 2004; Vázquez & Simberloff, 2002). Moreover, such anthropogenic food sources remain available throughout the year, thus providing a risky yet reliable and easily accessible resource option which is thought to be one of the major driving forces behind human-animal co-existence within urban settlements (Bateman & Fleming, 2012; Lowry et al., 2013; Thabethe & Downs, 2018; Widdows et al., 2015). In some cases, urban-dwelling free-ranging animals have been shown to acquire a preference toward anthropogenic food items to minimize their foraging activities, so that could invest more energy and time in nurturing social relationships which is essential to attain better fitness benefits (Bryson-Morrison et al., 2016, 2017; Hoffman & O’Riain, 2012; Saj et al., 1999; Sha & Hanya, 2013; Thatcher et al., 2019).

India has more than 400 mammalian species, of which 17 are non-human primates with different conservation status (Karanth et al., 2010; Honnavalli N. Kumara et al., 2010; Molur et.al.,2003) who have ecological as well as socio-cultural importance. Three of these non-human primate species i.e., Rhesus macaques (*Macaca mulatta*), bonnet macaques (*Macaca radiata*) and Hanuman langurs (*Semnopithecus entellus*) are frequently found in Indian cities, market places, temples, roadside settlements, etc., where they are often provisioned with human offered food items and space, worshipped and protected by *Hindus* (Sharma et al., 2011). Their wide distribution range and various feeding habits reflect their generalistic nature where they use a handful of novel strategies such as “coo-calls”, begging gestures, car raiding, etc. to acquire the available food items directly from humans (Arbib et al., 2008; Deshpande et al., 2018; Sha et al., 2009; Sinha, 2005). However, such close human-animal interaction is often lethal, affecting their chances of survival within urban ecosystems (Gosselink et al., 2007; Grinder & Krausman, 2001; Paul et al., 2016; Vijayan & Pati, 2002). Furthermore, these high-calorie urban food items could have a substantial effect on these animals’ physiology underlying their behavioural patterns, thereby reshaping intra and inter-specific group dynamics in contrast to their natural counterparts (Orams, 2002; Trave et al., 2017; Higginbottom & Scott, 2004). In this scenario, it is imperative to understand how the oppression of urban expansion has thinned down the natural resources and influenced the lives of these animals, leading to urban-adaptation in these species.

While several studies have been carried out on the naturally omnivorous macaques (Goldstein & Richard, 1989; Ganguly & Chauhan,2018; Laska, 2001; Oppenheimer 1977), to understand their opportunistic feeding behaviour to co-exist with human settlements, there has been no study yet to quantify and compare the dietary preference of folivorous Hanuman langurs in urban areas. Due to their deity value, this species is endowed with ample human provisioning. However, their specialized tripartite stomach structure largely aids in the digestion of a leafy diet (Bauchop & Martucci, 1968; Caton, 1999). Moreover, unlike the terrestrial macaques, the arboreal nature of Hanuman langurs (Khanal et al., 2018) is also barring them from availing enough human provisioning which could supplement their cellulose-based diet. Therefore, it seems to be all the more difficult for these large-bodied colobines to obtain sufficient resources that could be invested in maintaining reproductive fitness within an ecosystem where their natural food options are either unavailable or scarce to support their energy demand.

Grey langurs (*Semnopithecus entellus*), commonly called Hanuman langurs have colonized various parts of the Indian subcontinent, ranging from the desert to forest fringes, and have lived with a diversified resource structure and human interference (Ashalakshmi et al., 2014; Chetan et al., 2014; Oppenheimer,1977). In comparison to the other species of Hanuman langur, grey langurs’ (*Semnopithecus entellus*) social organization is highly flexible (Caton, 1999; Newton, 1988; Rajpurohit et al., 2006; Sterck, 1999) and is often modified by the male-male competition followed by infanticides (Broom et al., 2004; Hrdy SB 1974; Sharma et al., 2010). Besides unimale-bisexual troops, all-male bands are also common in these langurs (L. S. Rajpurohit et al., 2003). Even though they exploit a wide range of plant species including various plant parts, only a fraction of these so-called “Least Concern species” (IUCN 2003) can reach their reproductive age, which is again expected to reduce due to the adverse effect of urban encroachment (Kumara et.al., 2020). On the contrary, such urban settlements provide easy access to various anthropogenic low-fiber food sources which are mostly processed yet offer high-calorie to these ruminant folivores (Sayers, 2013). Few articles have reported human-langur cooperation through the food provisioning in Indian cities and towns, portraying them as ecological generalists in terms of habitat and diet. This mismatch in their expected and observed diet has made them one of the prime examples of Liem’s paradox which refers to the odd pairing of specialized anatomical features with a generalistic diet (Binning et al., 2009; Liem, 1980). However, it was later argued that such “asymmetry allows optimally foraging consumers to evolve phenotypic specializations on nonpreferred resources without compromising their ability to use preferred resources” (Robinson & Wilson, 1998). Hence, the development of alternate feeding patterns in these urbanized free-ranging gray langurs demands considerable scientific attention which could provide important insights into their eco-ethological adaptation for better management and policy-making to develop a sustainable urban ecosystem. Several studies have manifested ‘food-resources’ as one of the major contributing factors that limit primate’s group size and composition (Chapman, 1990). The Van Schaik model posits that in folivorous non-human primates, the intra-group scramble feeding-competition leads to differential reproductive success which has an immense role in establishing the hierarchical relationship within the group (Van Schaik 1989; Borries 1993). Therefore, the feeding behaviour of the grey langurs can provide interesting insights into the social dynamics, as well as the urban adaptation of the species. In this article, we have focused on the feeding preference of group-living, free-ranging grey langurs in the urban areas of West Bengal, India, in a habitat where they successfully co-exist with humans.

## METHODS

### Study area and study animals

Free-ranging langur troops were identified through regular census methods between September 2018 to December 2018 in various parts of West Bengal, India, of which three distinct langur troops (one in Dakshineswar (22.6573° N, 88.3624° E), one in Nangi (22.4973° N, 88.2214° E), and one in Sarenga, Nalpur (22.5307° N, 88.1840° E)) were selected for long-term field-based observations, considering the various level of human interferences received by the langurs (ESM1,2). Observers visited the areas at random times during the day and walked on all roads, by-lanes, and fields covered with vegetation of the above three selected urban settlements. Whenever a troop of langurs was sighted, it was observed *ad libitum* for a minimum of 15 minutes to a maximum of one hour, during which the observer recorded their location, troop size, and behaviours (later categorized as either intra- or inter-specific interactions). These data were used to categorize langurs into four distinct life stages based on their physical and behavioural characteristics (Infant-dark fur colour and fully dependent on adults for their movement and feeding; Juvenile-light fur colour similar to adults but smaller in size and partially dependent on adults for their movement and feeding; sub adult-fully independent of adults but yet to attain sexual maturity, body size is typically in between that of juveniles and adults; adult-fully grown, independent individual who is sexually mature) and also to note their territories (by using GPS-eTrex30, Garmin).

### Long-term study

For long-term field-based observations, we followed these three troops between January 2019 to March 2020 and recorded their behaviours using a combination of an equal number of instantaneous scans and all occurrences sessions (AOS) (Altmann, 1974). Whenever any feeding behaviour was observed, the observer recorded the details of the initiator, recipient, and their respective behaviours along with the food types (food-census) and location. Then, we categorized food items being eaten by the langurs into two distinct categories like ‘natural’ and ‘urban’ food sources (ESM 3a,b).

### Food-choice test

We carried out a choice-based experiment in a field set-up to find out the feeding preferences of free-ranging langurs in an urban settlement like Dakshineswar where they received maximum human interference, including maximum food offerings (ESM 3a) (manuscript in preparation) and depend mostly on ‘urban’ food sources, unlike the langurs of Nalpur and Nangi (Figure 1). We offered a food tray of cardboard, with four types of food items (which were recorded as the ‘most frequently eaten’ food items during the food-census), each of them having a comparable quantity and size (ESM 4), at any random times of a day between 0600 to 1800 hours. We used cauliflower and brinjal as ‘natural food’ items, whereas bread and peanuts were used as ‘urban food’ items. All of these offered food items were fresh and suitable for human consumption. These were presented on the food tray in random order to avoid any ‘side-bias’. Since peanuts were seen to be one of the most frequently eaten urban food items, we used it for the choice-based experiment and offered it in a small paper bag (which is usually used by people to offer peanuts to langurs during provisioning), making its quantity visually similar to the other food items. Based on the food-census data, we sub-divided the study area, Dakshineswar, into three distinct zones representing various feeding options available to langurs (ESM 5). The experimenter randomly chose one zone and presented the food tray to a spot where the maximum number of troop members can have equal access to the food tray. The experimenter either waited until the food tray was empty or waited for ten minutes if the food tray remained unattended or partially attended by the langurs, before closing the session. Once started, the experiment remained undisturbed i.e. no human interference was allowed and the entire experiment was video-recorded. In order to avoid any bias, which could influence subsequesnt trials, the videos were decoded only in April 2020, after the completion of all experiments. We conducted a total of 83 experiments in the field set-up of which 74 experiments (where the food trays were attended by the langurs without having any human interference) were considered for the final analysis (ESM 6).

**Figure 1:**
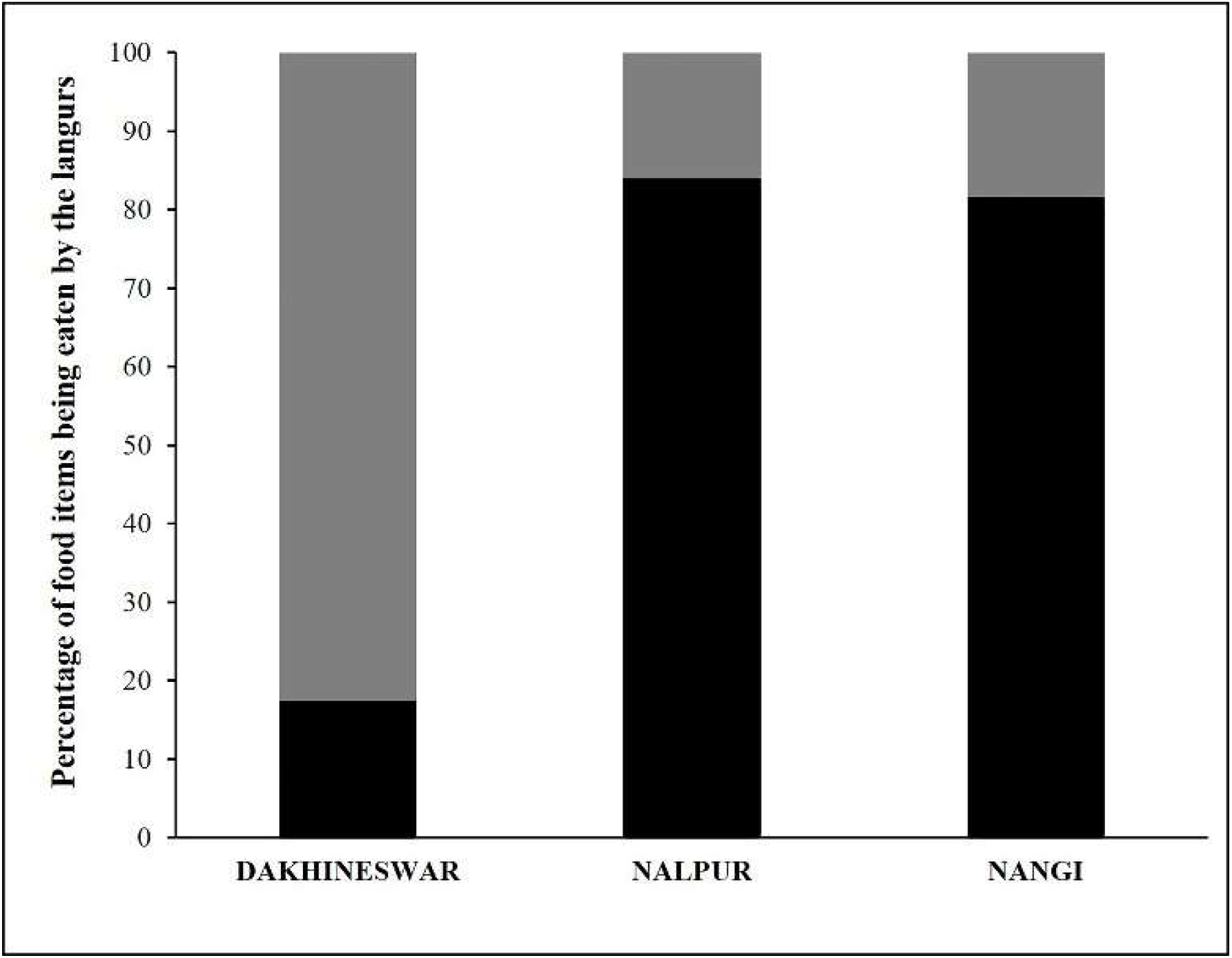
Stacked bar diagram showing the feeding habit of free-ranging langurs at three locations, Dakshineswar, Nalpur, and Nangi. Black and grey bars represent the percentage of ‘natural’ and ‘urban’ food items being eaten by the free-ranging langurs.

### Scoring method

For each experiment, we recorded the times (in seconds) when a food item was ‘approached’ or attempted to be received by the langurs (FA), chosen to be eaten (FC), the delay between FA and FC (in seconds) (delay), number of rejection received by a food item (RJ), and the presence or absence of aggression shown by the langurs to possess a food item (AG). Then we scored the food items for each of these categories to reflect the langurs’ feeding preference separately for each experiment. Since the food tray had four food items, each of them had five scoring options for FA and FC. The food that was attempted to be taken first received a score of five and the last (fourth) one was given a score of two. If a food item remained unattempted, it received a score of one. Similar scoring was done for FC, where the food scored ‘one’ if it was not chosen to be eaten and ‘five’ for being eaten first. RJ was scored on a scale of a maximum of ‘eight’ to a minimum of ‘zero’, where food items scored ‘zero’ if they were not rejected at all, and scored ‘eight’ when rejected for FA. Foods were scored ‘one’ if they received aggression and ‘zero’ if not, considering AG as an indication to the possessiveness for the most preferred food item which langurs did not want to share with. For ‘delay’ we scored them between ‘zero’ to ‘five’ where ‘zero’ represents no delay, ‘four’ for the maximum delay between FA and FC, and ‘five’ for the foods which were approached but not chosen to be eaten until the end of the experimental session.

### Statistical analysis

We used the scores for FA, FC, RJ, AG, and delay for all statistical analyses which were carried out using *StatistiXL* (*version 2.0*), and *R* (*version 4.0.2*). We ran a correlation analysis to check the inter-relation between various factors like ‘attempt’ (FA), ‘choice’ (FC), ‘delay’, ‘rejection’ (RJ), and ‘aggression’ (AG) which were affecting the final food selection by the langurs. To verify the results of the correlation we used a generalized linear models (GLM) and checked which parameter was finally affecting the final selection of the food items. We used the FC as the response variable, whereas FA, AG, RJ, and ‘delay’ were incorporated into the model as the predictor variables. We used a ‘Poisson’ distribution for the response variable to run the model. The distribution of the residuals was evaluated to check how well the model fits the data (ESM 7). A ‘principal component analysis’ (PCA) was conducted for descriptors like FA, FC, AG, RJ, and delay to check their effect on the food selection separately for three zones (Figure 2).

**Figure 2:**
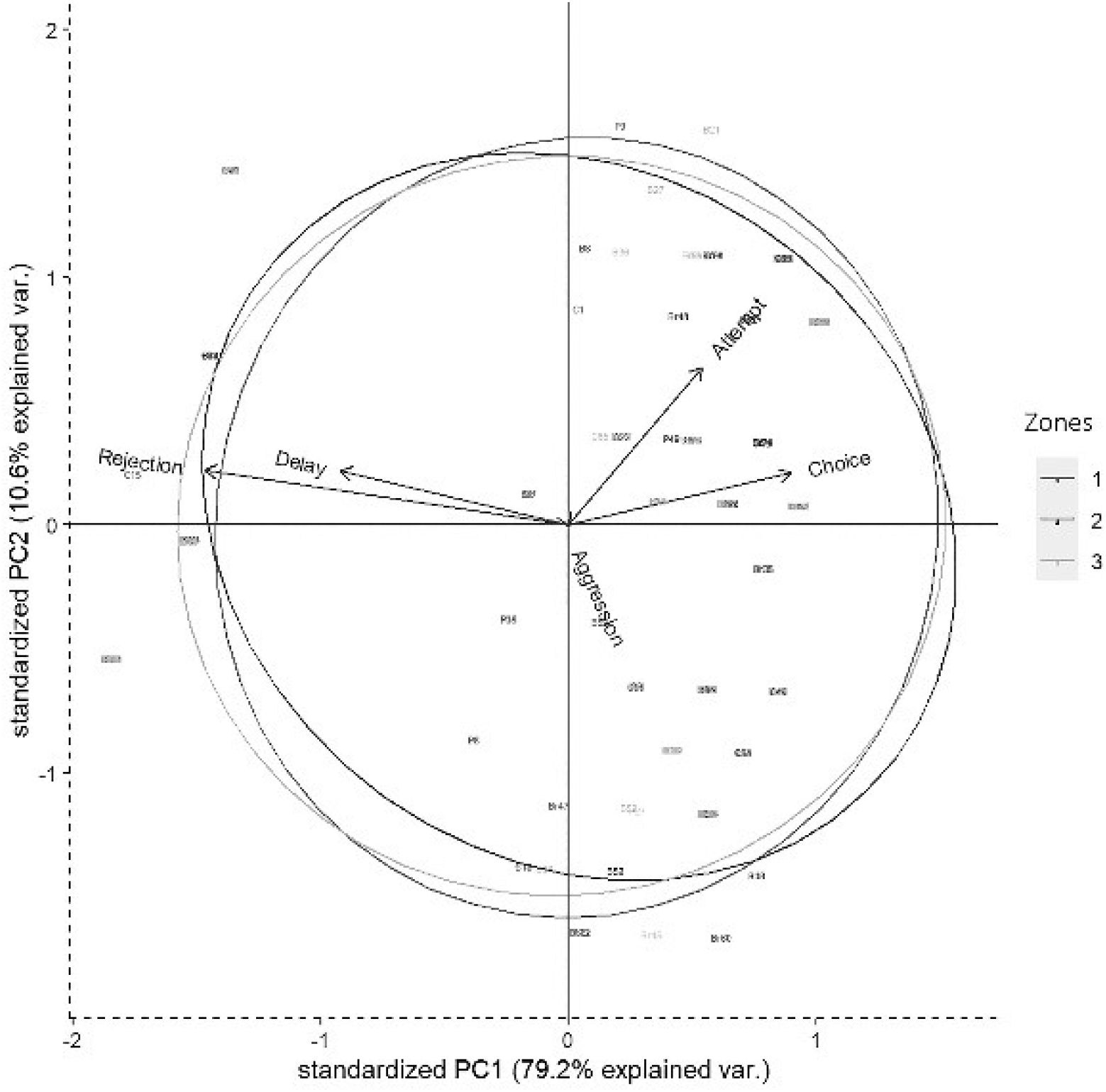
Biplot representing the distribution of variables in 2D space for the Principal component analysis (PCA) having descriptors like attempt, choice, delay, rejection, and aggression. Circles represent three different zones of Dakshineswar

### Multinomial logit model

To explain the preference of one food over another, i.e., food choice, we ran varied combination of multinomial logit models (MLM). Since we were interested in checking the predictive values of different independent variables like aggression, rejection, etc. on the final outcome of food choice, we ran 2 different sets of MLMs – separately for the ‘approach’ and ‘choice’ probabilities (Table 1). These two sets had four sub-models each where we employed a ‘leave one component out’ (LOCO) approach to meet our goal. The LOCO approach leaves one food component out at each sub-model step to check the order of selection of the subsequent food item. Besides, the models also evaluate the importance of the independent variables or descriptors in the outcome.

**Table 1:** Table representing the estimates, and p-values of the multinomial logit models, set 1 and 2 respectively for a) approach, and b) choice probabilities. These two sets represent four sub-models each. Here, we employed a ‘leave one component out’ (LOCO) approach which leaves one food component out at each sub-model step to check the order of selection of the subsequent food item.

| a) | Coefficients | Estimate | Std. Error | z value | p value |
| --- | --- | --- | --- | --- | --- |
| Bread is approached first | Attempt: Brinjal | 0.075 | 0.129 | 0.580 | 0.562 |
|  | Attempt: Cauliflower | -0.080 | 0.126 | -0.631 | 0.528 |
|  | Attempt: Peanuts | -0.293 | 0.126 | -2.322 | 0.020 * |
| Brinjal is approached first | Attempt: Bread | -0.075 | 0.129 | -0.580 | 0.562 |
|  | Attempt: Cauliflower | -0.155 | 0.128 | -1.206 | 0.228 |
|  | Attempt: Peanuts | -0.367 | 0.128 | -2.868 | 0.004 ** |
| Cauliflower is approached first | Attempt: Bread | 0.080 | 0.126 | 0.631 | 0.528 |
|  | Attempt: Brinjal | 0.155 | 0.128 | 1.206 | 0.228 |
|  | Attempt: Peanuts | -0.213 | 0.124 | -1.713 | 0.087 . |
| Peanuts is approached first | Attempt: Bread | 0.293 | 0.126 | 2.322 | 0.020 * |
|  | Attempt: Brinjal | 0.368 | 0.128 | 2.868 | 0.004 ** |
|  | Attempt: Cauliflower | 0.213 | 0.124 | 1.713 | 0.087 . |
| b) | Coefficients | Estimate | Std. Error | z value | p value |
| Bread is chosen first | Choice: Brinjal | -0.045 | 0.114 | -0.399 | 0.690 |
|  | Choice: Cauliflower | -0.309 | 0.113 | -2.747 | 0.006 ** |
|  | Choice: Peanuts | -0.374 | 0.113 | -3.301 | 0.001 *** |
| Brinjal is chosen first | Choice: Bread | 0.045 | 0.114 | 0.399 | 0.690 |
|  | Choice: Cauliflower | -0.264 | 0.111 | -2.371 | 0.018 * |
|  | Choice: Peanuts | -0.328 | 0.112 | -2.935 | 0.003 ** |
| Cauliflower is chosen first | Choice: Bread | 0.309 | 0.113 | 2.747 | 0.006 ** |
|  | Choice: Brinjal | 0.264 | 0.111 | 2.371 | 0.018 * |
|  | Choice: Peanuts | -0.065 | 0.109 | -0.596 | 0.551 |
| Peanuts is chosen first | Choice: Bread | 0.374 | 0.113 | 3.301 | 0.001 *** |
|  | Choice: Brinjal | 0.328 | 0.112 | 2.935 | 0.003 ** |
|  | Choice: Cauliflower | 0.065 | 0.109 | 0.596 | 0.551 |

The first set used the food approach as the outcome and the second set used the goal function of final food choice. When one food was approached i.e being attempted to be received by langurs, the probabilities of approaching the next food items can be determined subsequently by using set 1 MLMs. We ran four sub-set MLMs to check what would be the next approached food items separately while considering either brinjal, bread, cauliflower, or peanuts as the ‘first approached’ food item (Table 1a). Higher scores of the odds ratio confirm the results of MLM estimates thereafter (Table 2a) and subsequently rank the different food approach preferences according to the LOCO tactic.

**Table 2:** Table representing the odds ratio separately for a) approach, and b) choice probabilities.

| a) Approach probabilities |  |  | Odds ratio |
| --- | --- | --- | --- |
| Bread is approached first | Attempt: Brinjal |  | 1.078 |
|  | Attempt: Cauliflower |  | 0.923 |
|  | Attempt: Peanuts |  | 0.746 |
| Brinjal is approached first | Attempt: Bread |  | 0.928 |
|  | Attempt: Cauliflower |  | 0.857 |
|  | Attempt: Peanuts |  | 0.692 |
| Cauliflower is approached first | Attempt: Bread |  | 1.083 |
|  | Attempt: Brinjal |  | 1.167 |
|  | Attempt: Peanuts |  | 0.808 |
| Peanuts is approached first | Attempt: Bread |  | 1.340 |
|  | Attempt: Brinjal |  | 1.445 |
|  | Attempt: Cauliflower |  | 1.238 |
| b) Choice probabilities |  |  | Odds ratio |
| Bread is chosen first | Choice: Brinjal |  | 0.956 |
|  | Choice: Cauliflower |  | 0.734 |
|  | Choice: Peanuts |  | 0.688 |
| Brinjal is chosen first | Choice: Bread |  | 1.047 |
|  | Choice: Cauliflower |  | 0.768 |
|  | Choice: Peanuts |  | 0.720 |
| Cauliflower is chosen first | Choice: Bread |  | 1.362 |
|  | Choice: Brinjal |  | 1.302 |
|  | Choice: Peanuts |  | 0.937 |
| Peanuts is chosen first | Choice: Bread |  | 1.453 |
|  | Choice: Brinjal |  | 1.389 |
|  | Choice: Cauliflower |  | 1.067 |

Simultaneously, the set 2 MLMs were processed to establish and validate the preferred order of food items being chosen (final food choice) by the langurs during the experiment (Table 1b, 2b). The LOCO here assumes that one food has been consumed (and thus exhausted) and subsequently calculates the probabilities of choosing the next item. Since all food items were provided equally (i.e. equal probability of choice at the beginning), the model considered the frequencies of alternatives equal to 0.25. We used the Newton-Ralphson method from the package ‘*mlogit*’ in *R* to run the MLM (Croissant, 2020). The estimates of the MLM were plotted after normalizing to 1.0 (to avoid the negative values) for the visual representation (Figure 5).

### Food sharing

We recorded the incidents of food sharing between langurs, if any, out of the total 221 successful cases (where the food items were attended by langurs) from a total of 296 cases (four food options for 74 experiments). We included the details of the initiator, recipient, proportion of food being shared, and the interest of the initiator to share the food item with the recipient (i.e. shared forcefully or not). We used social network analysis (SNA) by using *Cytoscape* where we used various life stages (adult, subadult, juvenile, and infant) as a ‘node’ and an incident of food sharing between them as a ‘link’, separately for each food type (Figure 3). Here, we calculated the ‘*indegree*’ and ‘*outdegree*’ for each node representing the number of food sharing behaviour initiated and received by them respectively.

**Figure 3:**
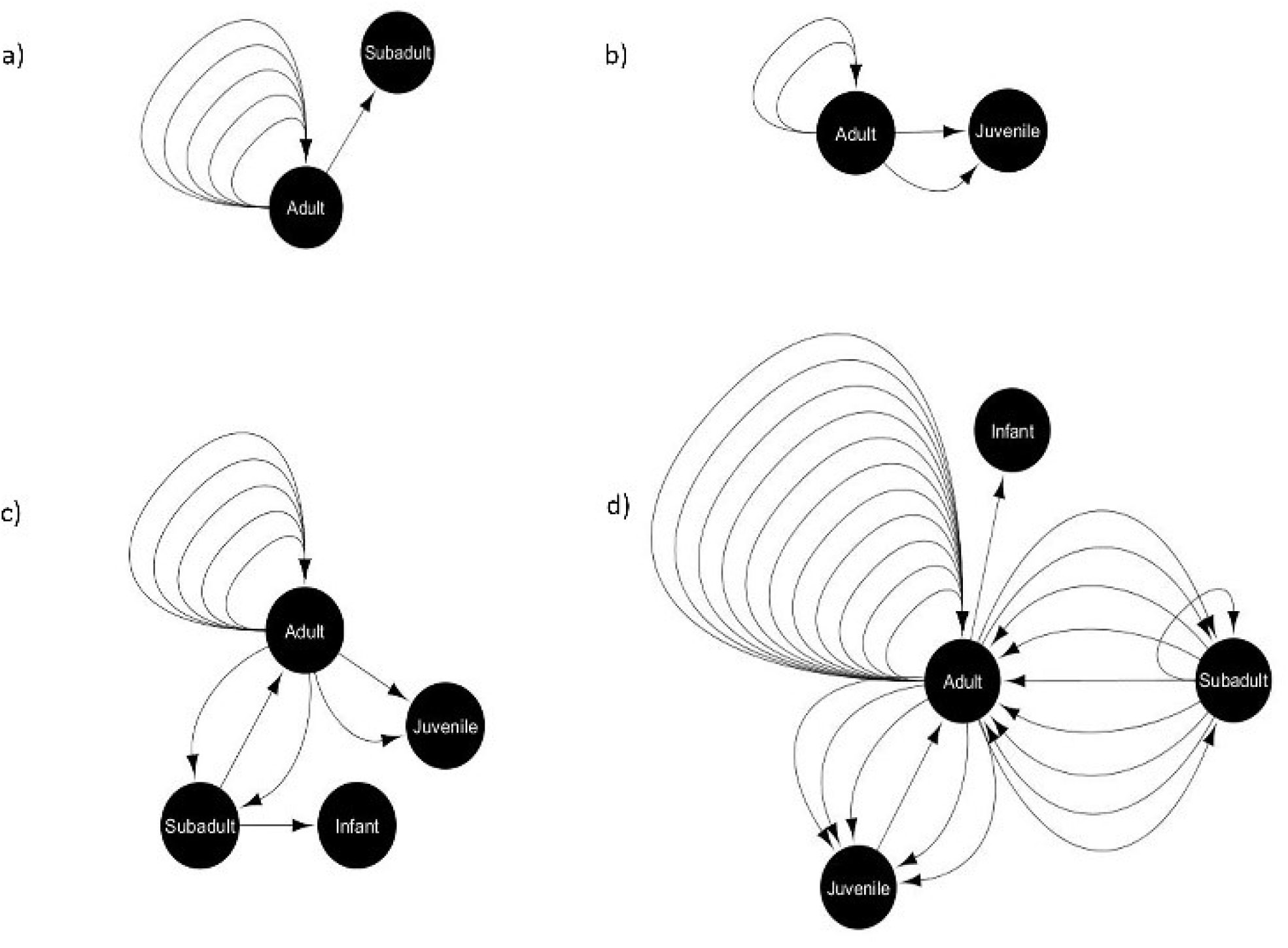
Food-sharing networks of free-ranging hanuman langurs for various food items like a) bread, b) brinjal, c) cauliflower, and d) peanuts. The solid black circle represents a node, depicting a particular life stage of langurs. The black arrow represents one food-sharing behaviour between two nodes, which originated from the initiator and directed towards the recipient.

### Ethical note

No langurs were harmed during this work. All work reported here was purely observation-based and did not involve direct handling of langurs in any manner, therefore, was in accordance with approved guidelines of animal rights regulations of the Government of India. The research reported in this paper was sanctioned by DST-INSPIRE, Government of India (approval number: DST/INSPIRE/04/2018/001287, dated 24^th^ July 2018), and was also notified to the Principal Chief Conservator of Forests (PCCF), West Bengal, India.

## RESULTS

### Food census

Feeding habit of free-ranging langurs greatly varied between locations (Contingency chi square: χ^2^ = 122.15, df = 2, p > 0.0001). Langurs of Dakshineswar largely depended on urban food sources (83%) which were mostly human offered (manuscript in preparation), whereas in Nalpur and Nangi they mostly relied on the natural food sources (84% and 82% respectively) (Figure 1).

### Food choice test

We carried out a total of 83 experimental trials in Dakshineshwar, of which 74 were successful. The experimental outcomes were perused by ‘Correlation analysis’ and ‘Generalized linear model (GLM)’. Correlation analysis-Rejection (RJ) was seen to be highly correlated to attempt (FA), choice (FC), and ‘delay’. A significant positive correlation (r = 0.755, p < 0.01) was found between RJ and ‘delay’. On the other hand, high negative correlations with FA and FC (r = FA: −0.504, FC: −0.814; p<0.01) represented inverse relations of the same with these factors. FC was highly and positively correlated to FA (r = 0.685, p < 0.01), especially towards a few food items like bread and brinjal (bread = 0.786, brinjal = 0.726, cauliflower = 0.594, peanuts = 0.606). On the contrary, ‘delay’ had significant negative effects on FC (r = −0.76, p < 0.01) (Figure 4).

**Figure 4:**
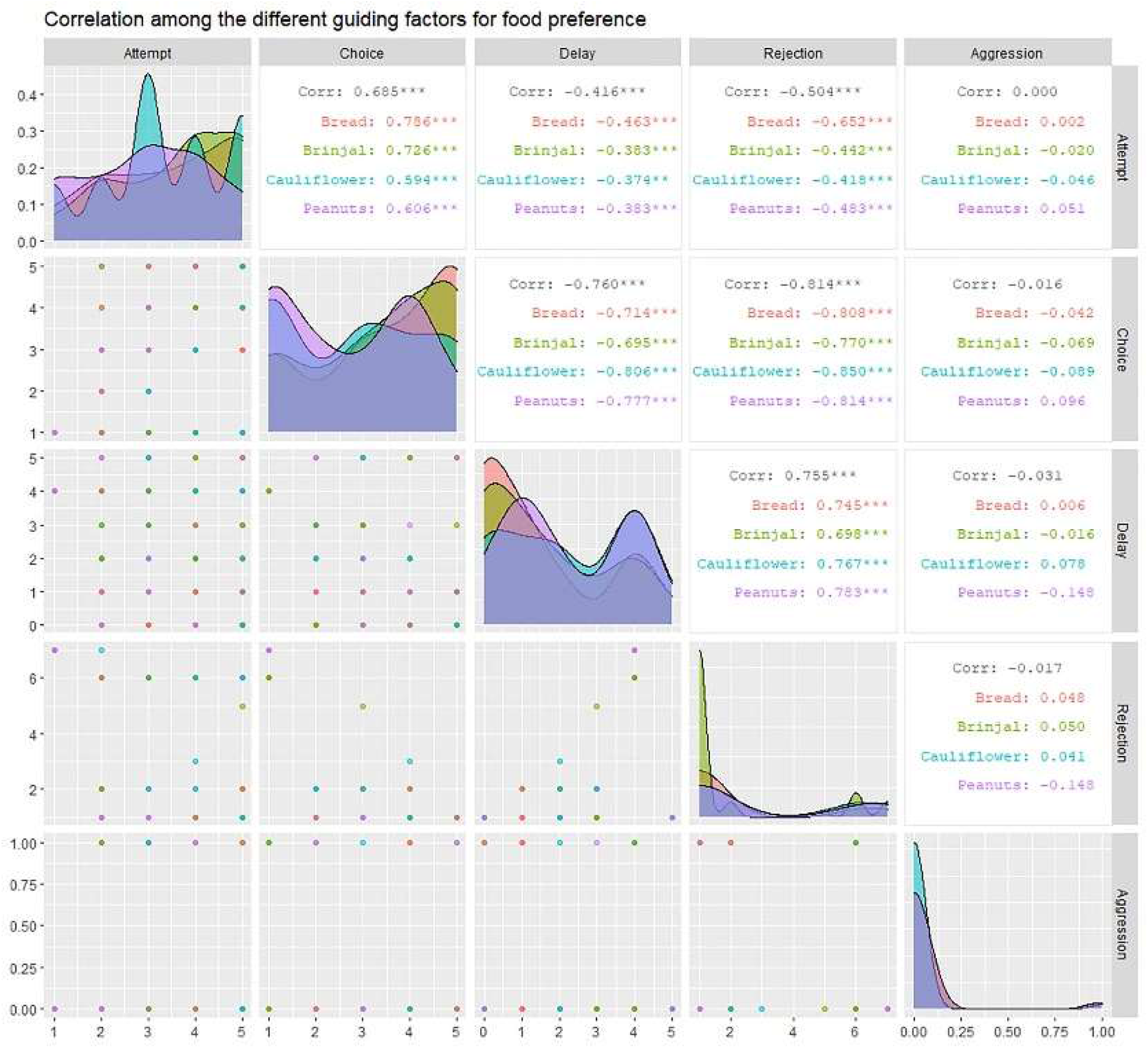
The correlogram representing the inter-relation between factors like ‘attempt’ (FA), ‘choice’ (FC), ‘rejection’ (RJ), ‘aggression’ (AG), and ‘delay’. It provides the correlation coefficient values (r) for each combination of factors and separately for each of the four food items along with their level of significance.

The GLM confirmed the significant effects of predictor variables like attempt, rejection and delay on the final food choice. Considering the estimates and p-values, while FA (positive) and RJ (negative) showed more significant effects on the FC (p < 0.01), ‘delay’ had a lesser impact (negative) (p < 0.05) (Table 3). An even distribution of the residuals on either side of ‘0.0 line’ indicated that the model had a good valid fit (ESM 7).

**Table 3:** Table showing the outcomes of the generalised linear model (GLM). ‘Attempt’ shows a positive estimate value for p < 0.01, whereas ‘rejection’ and ‘delay’ come up with negative estimate values for p < 0.01 and p < 0.05 respectively. Though ‘aggression has a negative estimate value, it is not significant. Significant codes: 0 ‘***’ 0.001 ‘**’ 0.01 ‘*’ 0.05 ‘.’ 0.1 ‘.’ 1.

|  | Estimate | Std. Error | z value | p value |
| --- | --- | --- | --- | --- |
| Attempt | 0.152 | 0.031 | 4.950 | 7.43e-07 *** |
| Rejection | -0.171 | 0.028 | -6.102 | 1.05e-09 *** |
| Delay | -0.067 | 0.030 | -2.252 | 0.0243 * |
| Aggression | -0.041 | 0.158 | -0.259 | 0.7957 |

Aggression (AG) had a slight negative influence on both FC and RJ (−0.29 ≤ r ≤ 0 i.e. weak negative) (Figure 4, Table 3). However, the linear model (LM) plot revealed that when AG was not present (left panel, aggression = 0) and ‘delay’ was minimum (red color bands), FC was highest for lower RJ and *vice-versa*. Right panel showed that the presence of AG increased the ‘delay’ in FC (width of the colour bands represents the ‘increase’) (Figure 6a). Furthermore, a detailed LM plot revealed that with an increase in the FA, probabilities for FC increased, but both RJ and ‘delay’ lowered the FC (Figure 6b).

**Figure 5:**
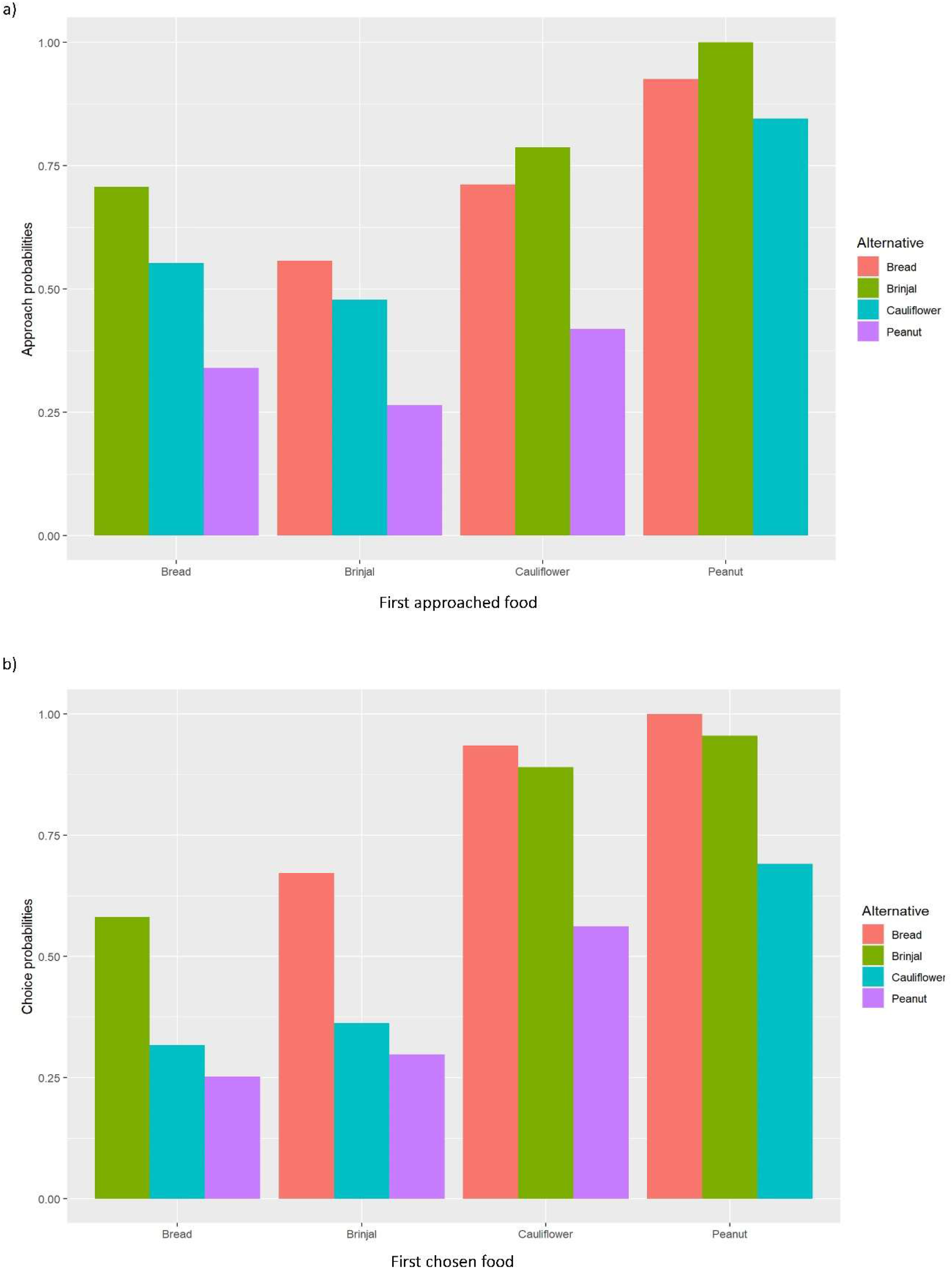
Bar diagrams representing the normalized values of MLM estimates separately for a) ‘approach’, and b) ‘choice’ probabilities. The X-axis represents the a) first approached and b) first chosen foods.

**Figure 6:**
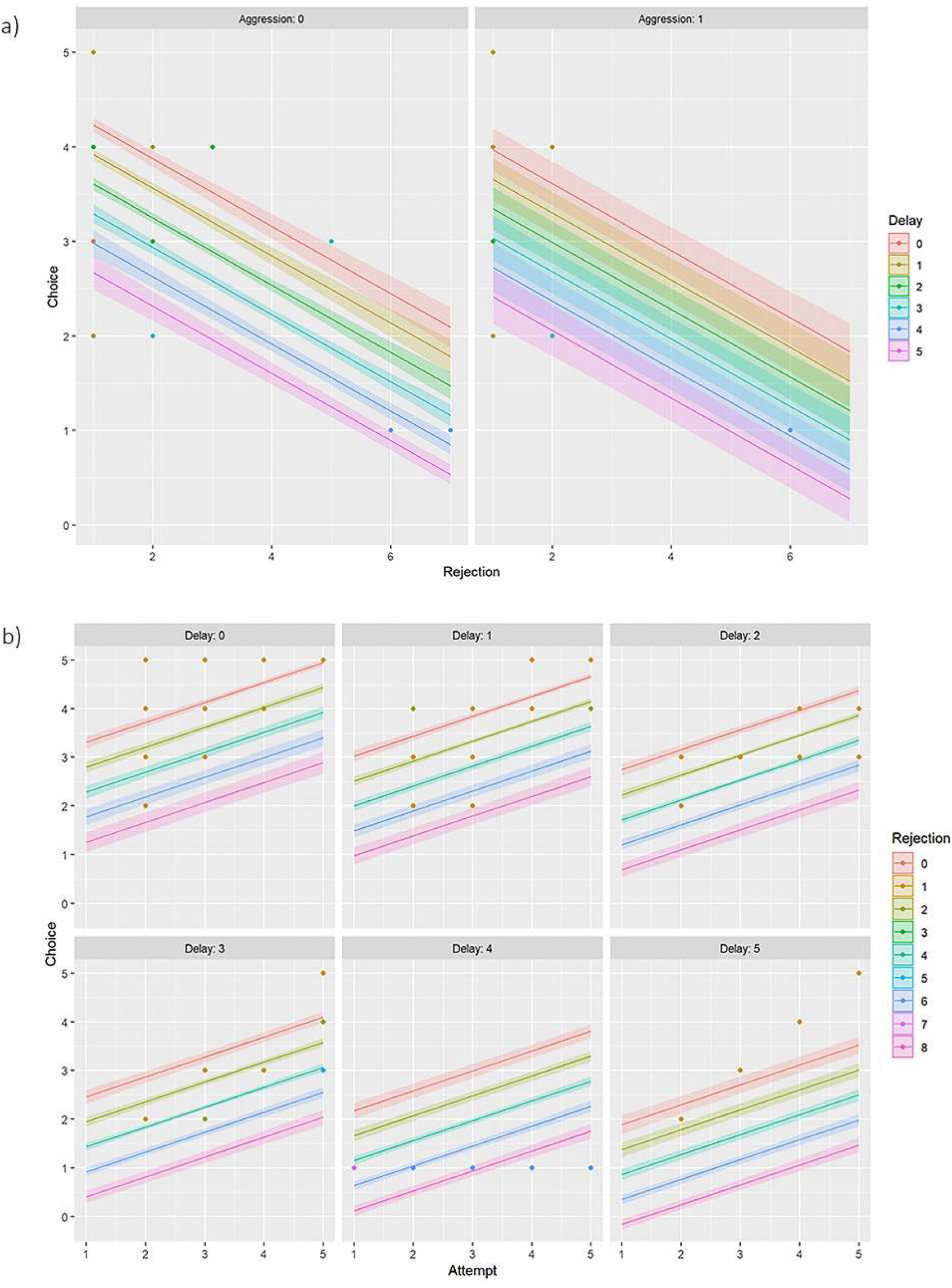
Linear model (LM) plot representing the variations in the choice of food for ‘rejection’ and ‘delay’. The width of the colour bands increases with the delay. a) LM plots showing different levels of ‘aggression’ has different effects on food choice. The left panel represents data for ‘zero aggression’, whereas the right panel shows that the ‘presence of aggression’ increases the delay in food choice. b) LM plots showing the effects of ‘attempt’ on food choice, together with ‘delay’ and ‘rejection’. Each panel represents a particular ‘delay’ score. For example, the top left panel is for ‘no delay’ or ‘zero’ delay score, and the bottom right is for the ‘maximum delay’ i.e. score five.

### Principal component analysis (PCA)

Results of PCA showed that most of the variability in the experimental observations could be explained through PC1 (79.20%), and subsequently another 10.60% by PC2 (Table 4). The PCA biplot revealed that the ‘zones’ had no impact on food selection by langurs. The arrows associated with descriptors ‘attempt’ and ‘choice’ remained close to each other, and pointed in the direction of the increasing values of both PC1 and PC2 (the signs of the eigenvectors are also positive for both PC1 and 2, Table 4, Figure 2), thereby confirming their positive effects on the food selection. However, ‘delay’ almost overlapped with the ‘rejection’ and pointed in the direction of the low value of PC1 but high value of PC2 (the signs of the eigenvectors for PC1 is negative and positive for PC2, Table 4, Figure 2), revealing their negative impact on the food selection. The individual loading of ‘aggression’ was only −0.99 on PC5, therefore considered to have a minimal effect (Table 4).

**Table 4:** Tabulated representation for the principal component analysis (PCA). It represents the ‘proportion of variance’ for each principal component (PC), followed by a ‘cumulative proportion’. It also displays the individual loading for each descriptor like ‘attempt’, choice’, ‘delay’, ‘rejection’, and ‘aggression’.

|  | PC1 | PC2 | PC3 | PC4 | PC5 |
| --- | --- | --- | --- | --- | --- |
| Proportion of Variance | 0.792 | 0.106 | 0.072 | 0.026 | 0.004 |
| Cumulative proportion | 0.792 | 0.898 | 0.970 | 0.996 | 1.000 |
| Loadings of eigenvectors: |  |  |  |  |  |
| Attempt | 0.266 | 0.859 |  | 0.437 |  |
| Choice | 0.444 | 0.285 | 0.153 | -0.835 |  |
| Delay | -0.456 | 0.298 | -0.789 | -0.284 |  |
| Rejection | -0.724 | 0.303 | 0.595 | -0.172 |  |
| Aggression |  |  |  |  | -0.999 |

### Multinomial logit model

The multinomial logit model (MLM) provided a higher score for bread (estimate value: −0.075) among others, revealing the probability of approaching bread as the 2^nd^ alternative, followed by cauliflower (estimate value: −0.155), and peanuts (estimate value: −0.368) when brinjal was attempted first. Similarly, the MLM picked up bread, cauliflower, and peanuts, one by one, as the first attempted food item, and checked the probability of approach for the rest. Together with MLM estimate values, odds ratio confirmed the highest approach probabilities for brinjal, followed by bread, cauliflower, and peanuts (Table 1a, 2a, Figure 5a). However, for the choice probabilities, bread scored highest for both the MLM estimates and odds ratio, followed by brinjal, cauliflower, and peanuts (Table 1b, 2b, Figure 5b).

### First attempted vs eaten food

Bread and brinjal were chosen as the first attempted food item (scored ‘five’ for FA) for 31% and 32% cases respectively, followed by cauliflower (23%) and peanuts (14%). However, not always the first attempted foods were chosen to be eaten first. Langurs switched their preference between the first attempt to first choice for 29.7% cases, and mostly for bread (Goodness of fit: χ^2^ = 31.08, df = 3, p<0.0001) (ESM 6).

### Food sharing

Only 18% of the total successful cases were recorded where langurs shared the received food items with their troop members during the experimental trials. However, the shared food items mostly consisted of the least preferred peanuts (53%), and cauliflower (22%) (Goodness of fit: χ^2^ = 44.72, df = 3, p < 0.0001) (ESM 8). Social network analysis revealed that food sharing mostly occurs between adults (Goodness of fit: *Outdegree:* χ^2^ = 75.35, df = 3, p < 0.0001; *Indegree:* χ^2^ = 40.39, df = 3, p < 0.0001) and largely for peanuts, and cauliflower (Figure 3).

## DISCUSSION

Folivores colobines have received considerable research attention because of their unique ability to ingest large quantities of foliage (Newton, 1992; Oates 1988; Struhsaker & Oates, 1975). Their multipartite stomachs are lined with mucus-secreting glands which facilitate the fermentation of leafy diet in the presence of cellulolytic bacteria (Caton, 1999). However, the dietary composition of free-ranging Hanuman langurs (*Semnopithecus entellus*) seems to be relatively complex. They often use a diverse array of plant parts including leaves, stalks, shoots, buds, flowers, and fruit to utilize the available resources at its best (Yoshiba, 1967; Vandercone et al., 2012). Besides, Srivastava and Winkler added insectivory and human-provisioning to the Hanuman langurs’ feeding repertoire (Srivastava, 1989,1991; Winkler 1988). However, these feeding habits were mostly seasonal and plant parts still accounted for a significant portion of their regular diet (Koenig & Borries, 2001) similar to the langur group of Nungi, and Nalpur.Surprisingly, the langur troop of Dakhineswar, West Bengal, India, was spotted to thrive largely (83% of the total diet) on the ‘urban’ food sources for their sustenance within human settlements, throughout the year. Similar to other free-ranging scavengers like dogs, jackals, monkeys, (Butler & du Toit, 2002; Paul et al., 2016; Sanyal et al., 2010) this langur troop was observed to rely upon human generosity and food provisioning, seeking easy access to the ‘urban’ food sources (Dasgupta et al., manuscript in preparation). However, unlike these carnivorous, and omnivorous scavengers, langur’s stomach physiology looks alike to that of herbivores such as Macropodidae (Caton, 1999). Therefore, human-provisioned ‘urban’ food sources could have an inevitable health impact, followed by potential behavioural alteration in these urban-adapted free-ranging langurs.

Our field-based observational data reflected the highest degree of human interference in Dakhineswar where langurs frequently approached humans to acquire ‘urban’ food sources, in contrast to the langurs of Nangi, and Nalpur where they opted for foraging and scavenging and depended mostly on ‘natural’ food sources. Therefore, high human-langur interactions could be considered as an intriguing driving force behind langurs’ altered feeding habits in Dakhineswar. Moreover, the scarcity of plants and crop fields within the urban settlement like Dakhineswar might be another reason behind langurs’ consistent dependence on the ‘urban’ food options. In this context, the choice-based field experiment allowed us to understand whether it was the scarcity of the ‘natural food’ sources or the easy accessibility of the ‘urban food’ sources that lured them to get accustomed to the urban ecosystem.

The experimental set up allowed langurs to choose between natural and urban food sources, keeping aside the factors like scantiness, and human influences. Langurs chose brinjal, and bread consistently either as the first or second food options, in all the three zones of Dakhineswar, reflecting their feeding preferences within urban settlements. The outcomes of the experiments manifested the ‘attempt’ to be a significant precursor to the food selection. Moreover, its close association to the descriptor, ‘food choice’, for the increasing values of both principal components 1 and 2 in the PCA confirmed the significance of langur’s approach towards a particular food item. Therefore, it can be interpreted that the food had to be approached first prior to the final selection, allowing langurs for choice-based decision-making. However, the effects of ‘rejection’ and ‘delay’ were also substantial, and the final food selection by the langurs seemed to be non-random but a consequence of all the above three factors. A significant positive correlation between ‘rejection’ and ‘delay’ revealed that more delays in food selection might lead to the ultimate rejection of that particular food item. A greater rate of rejection negatively facilitated the final choice, whereas swifter attempts towards food led to less rejection. Therefore, when an increased ‘attempt’ escalated the probabilities for the final food selection, ‘delay’ gave rise to a dilemma between ‘food choice’ and ‘rejection’ which finally lowered the chances of selection for a given food item. ‘Aggression’ also had some negative effects on both ‘choice’ and ‘rejection’. Although it increased the ‘delay’ in final food selection, langurs used aggression to possess their chosen food items without being forced to share it with other troop members. Hence, our experiments revealed an active preference-based food selection by langurs within an urban settlement like Dakhineswar, driven neither by human interference, nor the scantiness of natural food options but by a keen interest towards specific food items. The multinomial logit model contemplated all of these factors for the final food selection by the langurs and revealed brinjal and bread to be the most attempted and first approached food items, followed by cauliflower and peanuts. However, bread outperformed brinjal as the most chosen food item for which langurs often switched their first approached food to the final selection, indicating their inclination to the urban food option.

In the case of food-provisioning where humans provide a food item of their choice to the animals, the animals have no scope to choose but to receive the offered food items. In our experimental set-up langurs had the liberty to choose from a platter of offered food items, without any human interference, and the underlying assumption was that the outcome of the experiment would be influenced only by their preference, if any. Our findings suggested that these langurs engaged with the items that were offered in the food-tray and chose bread and brinjal solely based on their feeding preference. Moreover, the social network analysis revealed that the adult langurs occasionally had the privilege to receive a food share from the focal langur, which rarely comprised of bread or brinjal, in contrast to the juveniles and subadults who hardly managed to get access to it, thereby showing their fondness for it.

### Conclusions

Although the impact of such ‘urban’ food sources on Hanuman langurs’ physiology is still debatable (Geffroy et al., 2017; Maréchal et al., 2016), it can be interpreted that these free-ranging Hanuman langurs of Dakhineswar not only learned to approach human-provisioned ‘urban’ foods but they acquired preferences for some of it which could facilitate their co-existence within an urban ecosystem. However, resource provisioning is often being correlated to peoples’ intension to get in touch with the wildlife, imposing a considerable threat to free-ranging animals’ survival chances (Orams, 2002; Trave et al., 2017). Yet, such man-animal interaction opens up possibilities for alternative easy access to resources like food and shelter for these animals who have lost their home due to urban encroachment (Cox & Gaston, 2018; Lowry et al., 2013; Theobald et al., 1997). Moreover, it has been shown that the ability to digest carbohydrates provided ancestral dog populations an advantage over wolves, facilitating the process of domestication, as the dogs could now utilize human-generated resources (Axelsson et al., 2013). Undoubtedly, our experimental results are an example of such urban adaptation where folivorous arboreal Hanuman langurs find their interest in terrestrial urban food items. Therefore, besides their deity value, the free-ranging Hanuman langurs’ successful co-existence with humans and their wide distribution throughout the Indian subcontinent could be well-explained by their altered yet opportunistic feeding pattern.

## Supporting information

ESM 6

## ACKNOWLEDGEMENTS

DD and AB equally contributed to this work. DD, RK, DB, SM, SK, ABh, and SG carried out the field work. MP and DD coded the entire data. AB and MP carried out all the statistical analyses. AB prepared the correlation matrix, ran PCA, GLM, and MLM to interpret the data. MP conceptualized the study, got grants to support the work, designed the fieldwork and supervised the work. MP, DD, PB drafted the manuscript.PS prepared the GIS map. PB provided the laboratory support to carry out the analysis. Author AB acknowledges the DSKPD Fellowship of University grants commission (no. BL/17-18/0490). All the authors acknowledge Dr. Anindita Bhadra, Associate Professor, Indian Institute of Science Education and Research-Kolkata, India, for her valuable inputs to the study and gave final approval for publication.

