## Supplementary Information for "Altered food habits? Understanding the feeding preference of free-ranging Grey langur (*Semnopithecus entellus*) within an urban settlement"

**Legends to Supplementary materials**

**ESM 1:** Map showing the three study locations

**ESM 2:** Table showing the group composition of three langur troops of Dakshineswar, Nangi, and Nalpur.

**ESM 3a:** Table showing various types of ‘natural’ and ‘urban’ food items being eaten by the free-ranging langurs of Dakshineswar, Nangi, and Nalpur.

**ESM 3b:** Table shows the calorie values of natural and urban food items commonly available in the Dakshineswar.

**ESM 4:** A ‘pitchboard’ made food-tray having four food items (from top right corner- bread, cauliflower, brinjal, and peanuts) being used for the choice-based field experiment.

**ESM 5:** Table representing three zones of Dakshineswar. Langurs were observed to depend mostly on human offered food items in ‘zone 1’, whereas in ‘zone 2’ and ‘zone 3’ they depend on both human-offered food items and food items being collected through foraging or scavenging. However, in ‘zone 2’ the foods largely consisted of foraged or scavenged items, but human offered items in ‘zone 3’.

**ESM 6:** Video of food choice experiment

**ESM 7:** A residual plot for the generalized linear fit model.

**ESM 8:** a) Pie chart representing the percentage of food items being shared between langurs out of the total successful cases. Larger pie on the left side represents the total 296 options of which 221 were successful (black coloured section of the pie). Smaller pie represents the 221 successful cases where the black dotted patch represents cases where the food sharing was observed. b) Pie chart representing the percentage of total food sharing separately for four food items

**Supplementary Information**

**ESM 1**

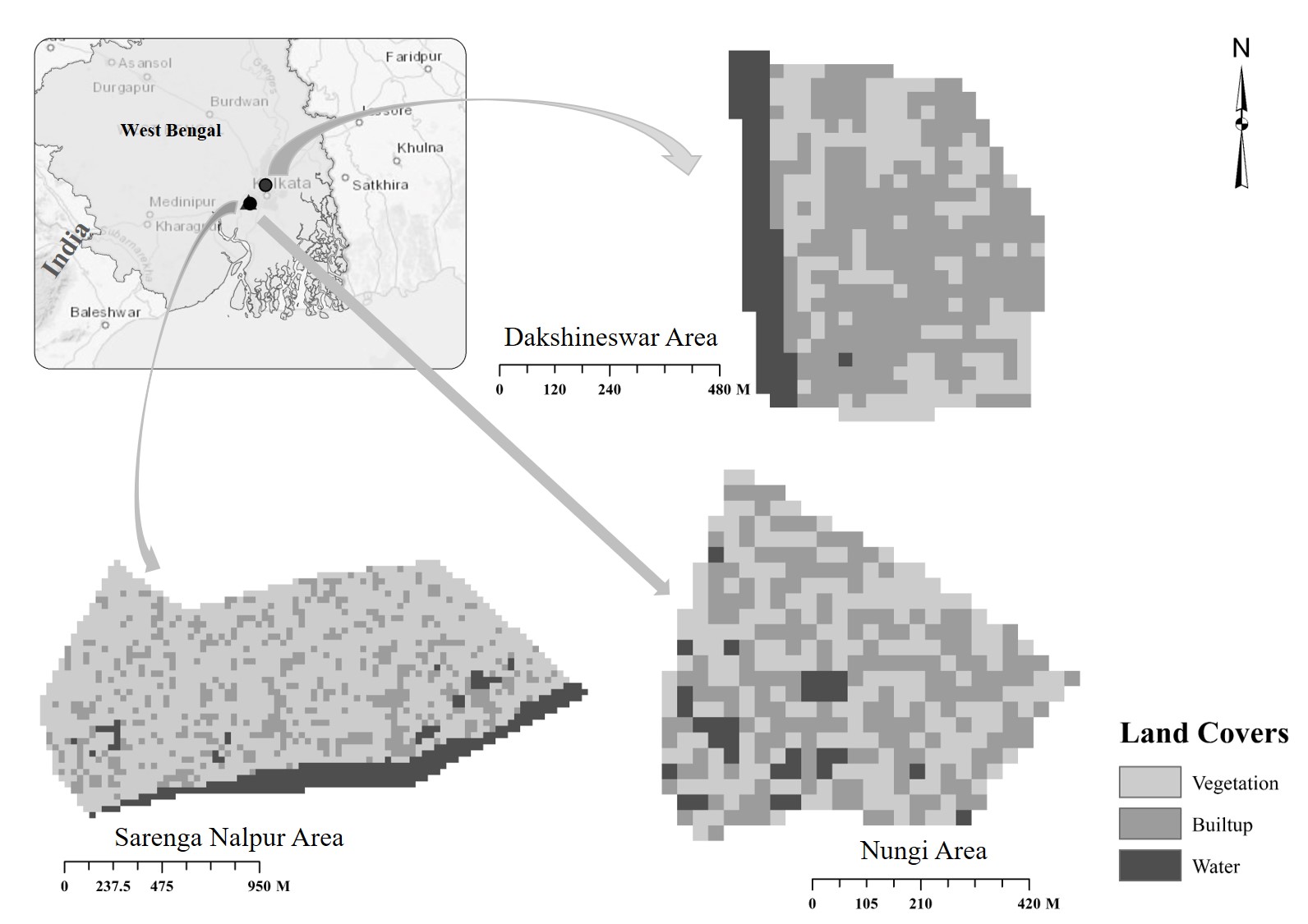

**ESM 2**

| Location | Adult | | Subadult | Juvenile | Infant | Total | Human interferences received  (Average frequency/hr.) |
| --- | --- | --- | --- | --- | --- | --- | --- |
|  | Male | Female |  |  |  |  |  |
| Dakshineswar | 2 | 8 | 2 | 4 | 2 | 18 | Highest (8/hr.) |
| Nangi | 1 | 5 | 3 | 3 | 1 | 13 | Medium (3.8/hr.) |
| Nalpur | 1 | 8 | 5 | 3 | 1 | 18 | Lowest (1.1/hr.) |

**ESM 3a**

| Sl. No. | Natural food items | Sl. No. | Urban food items |
| --- | --- | --- | --- |
| 1. | Leaves | 1. | Bread |
| 2. | Seeds | 2. | Peanuts |
| 3. | Fruits (both from the tree and market) | 3. | Chips |
| 4. | Vegetables (both from the crop field and market) | 4. | Cake |
| 5. | Tender stems | 5. | Icecream |
| 6. | Roots | 6. | Fried foods |
| 7. | Flowers | 7. | Puffed rice |

**ESM 3b**

| **Food category** | **Sl. No.** | **Food type** | **Calorie**  **(per 100 grams)** |
| --- | --- | --- | --- |
| Natural | 1. | Leaves | 15 |
|  | 2. | Tomato | 18 |
|  | 3. | Brinjal | 25 |
|  | 4. | Cauliflower | 25 |
|  | 5. | Mango | 60 |
|  | 6. | Guava | 68 |
|  | 7. | Potato | 77 |
|  | 8. | Banana | 89 |
| Urban | 9. | Bun | 300 |
|  | 10. | Peanuts | 567 |

References:

<http://www.fitbit.com/foods/1+bun/170624>

<http://www.fitbit.com/foods/food?viewFood=on&foodId=698398372>

<https://www.nutritionix.com/food/cauliflower/100-g>

<https://www.nutritionvalue.org/Eggplant%2C_raw_nutritional_value.html>

**ESM 4**

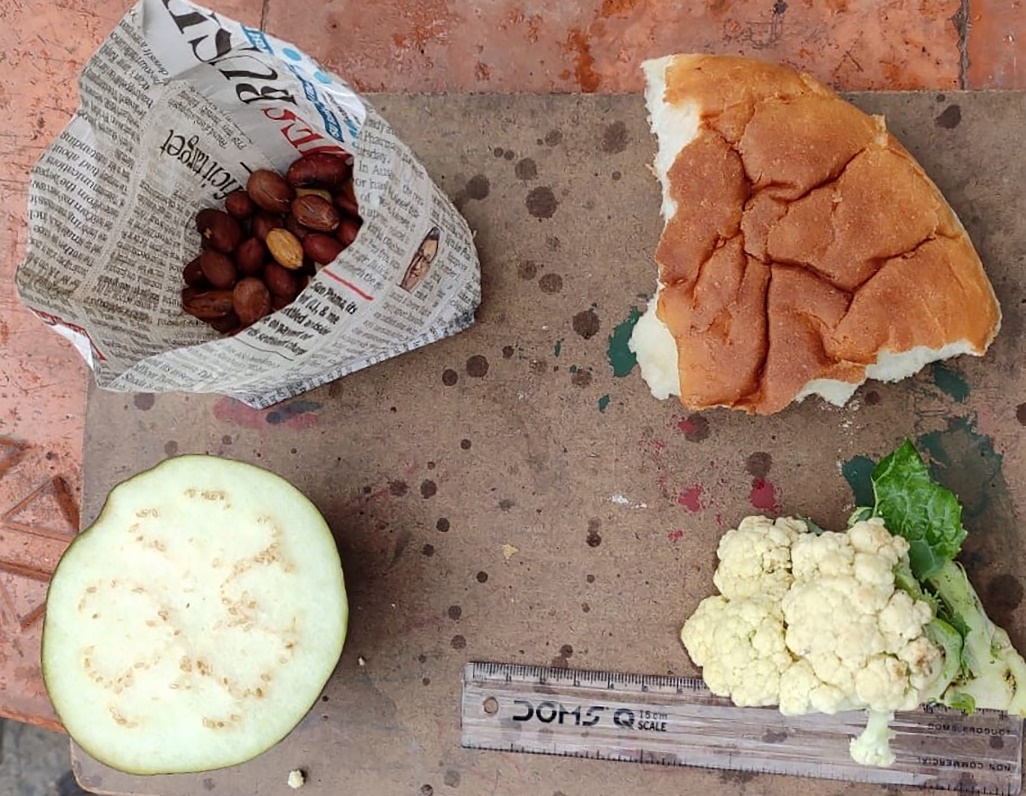

**ESM 5**

| Zones | Feeding options available to free-ranging langurs |
| --- | --- |
| 1 | Human offerings only |
| 2 | Foraging and scavenging > Human offerings |
| 3 | Human offerings > Foraging and scavenging |

**ESM 6**

See the video file named “Dasgupta et al. ESM6”.

**ESM 7**

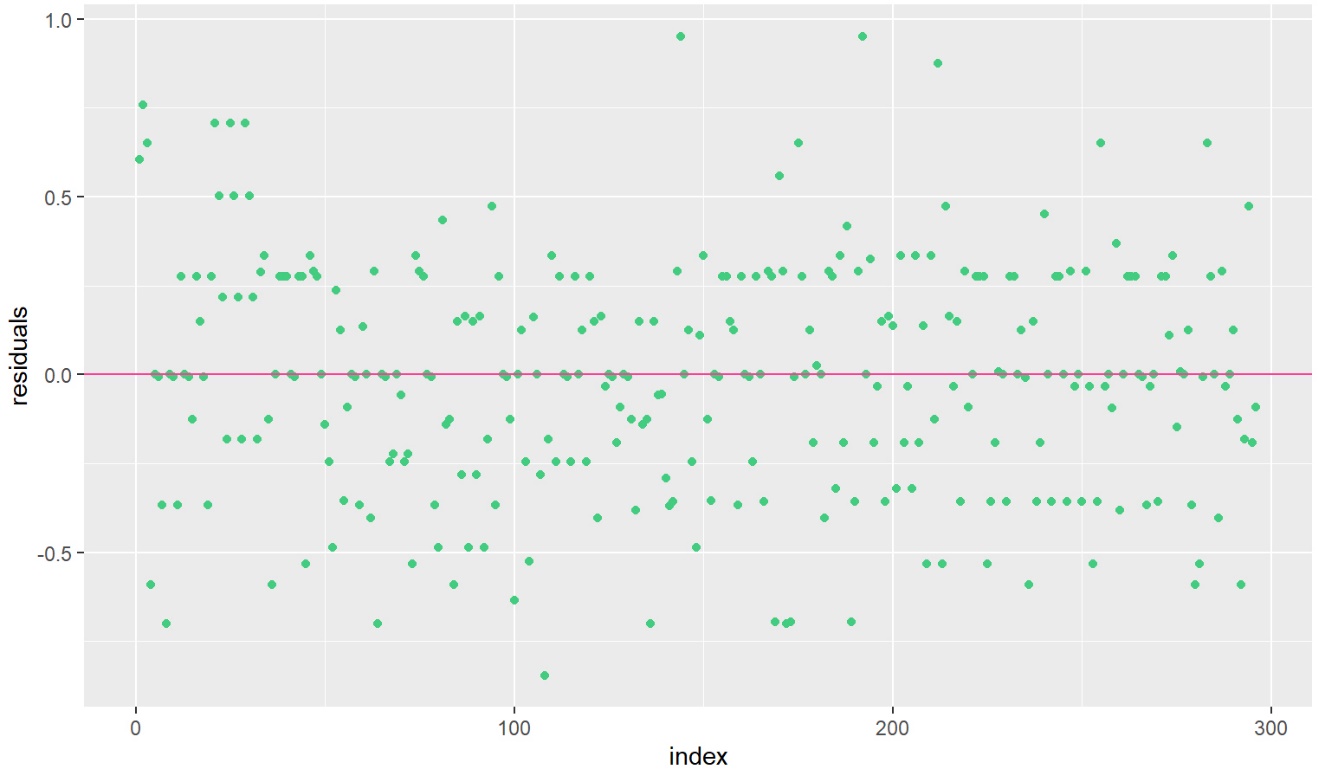

**ESM 8**

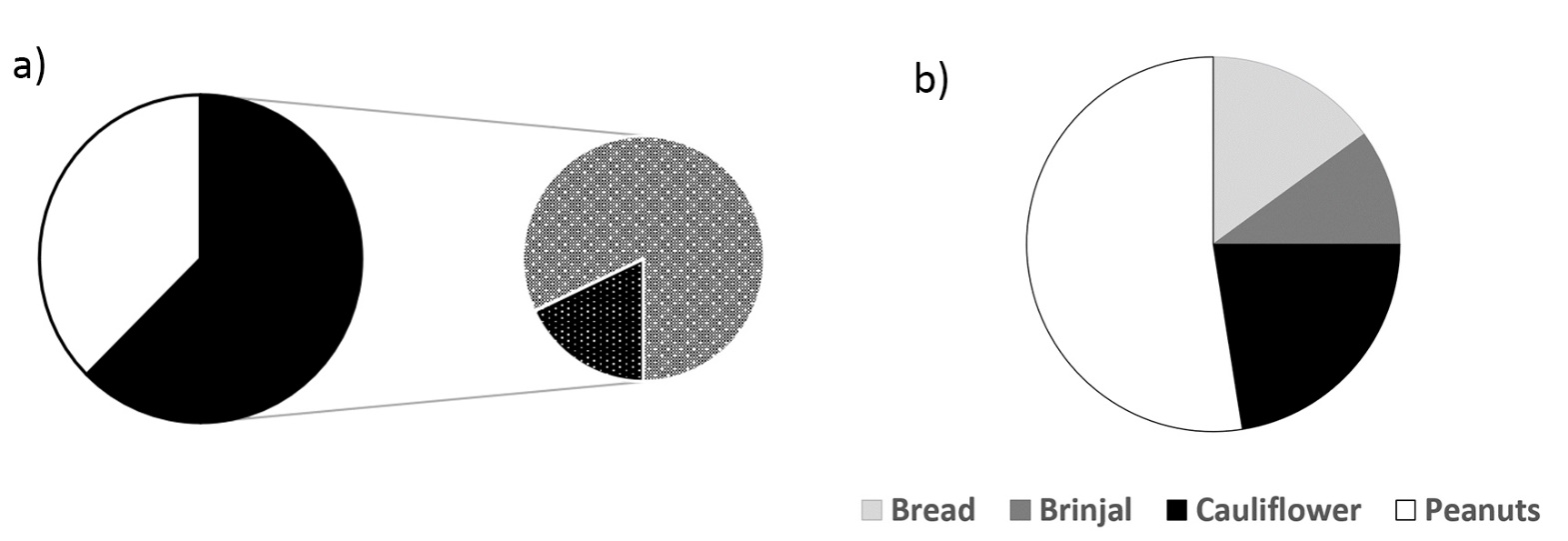
